# Manipulating plant development by editing histone methylation with the dCas9 tool: the *CUC3* boundary gene as a case study

**DOI:** 10.1101/2024.03.18.585636

**Authors:** Kateryna Fal, Marie Le Masson, Alexandre Berr, Cristel C. Carles

## Abstract

Chromatin modifications are deemed to associate with gene expression patterns, yet their causal function on transcription and cell fate remains unestablished. Here, we demonstrate the direct impact of an epigenome editing tool designed to remove a key chromatin modification at a precise locus in living plants, with outcomes from the molecular to the developmental scale.

The manipulated mark, H3K27me3, deposited at Lysine 27 of Histone 3 by the methyltransferase Polycomb PRC2 complex, is associated with the repression of developmental genes. As a new approach to investigate this histone mark genuine function, we used a dCas9-derived tool to bring a specific demethylase function at the *CUP SHAPED COTYLEDON 3* (*CUC3)* organ frontier gene, aiming to remove the trimethyl mark at H3K27. We show that the removal of H3K27me3 at the locus causally induces activation of *CUC3* expression within its regular territory, as well as ectopically. Our precise perturbation strategy reveals that alterations in a chromatin mark lead to changes in transcription and developmental gene expression patterning, with sharp consequences on plant morphogenesis and growth.

Our work thus constitutes a proof of concept for the effective use of epigenome editing tools in unveiling the causal role of mark dynamics, supported by both molecular and developmental evidences.

## Results and discussion

Considerable progresses have been achieved in uncovering the genetic and epigenetic regulators of development in multicellular eukaryotes. Among them, key players are the chromatin complexes that bring post-translational modifications on histone tails and modulate access to DNA for the transcriptional machinery^1–3^. In particular, the trimethyl mark deposited at Lysine 27 of Histone 3 (H3K27me3) is considered to control the dynamic regulation of key developmental genes, defining their spatial and temporal expression patterns and ensuring correct body plan establishment^4–6^. This role for H3K27me3 has largely been deduced from characterization of loss-of-function mutants in writers/erasers/readers, as well as from genome-wide profiling of marks and factor binding at the chromatin. Yet, such approaches are intricate due to multifaceted interactions of the chromatin mark propagators, including their activity on non-histone substrates, their non-catalytic functions, and the functional specialisation or redundancy of regulators within a same family, especially in plants^4,7–9,10^. For these reasons, indirect functional studies allowed drawing only limited and correlative conclusions on the relationships between H3K27me3 marks, transcriptional activity, gene expression and body plan organization.

Therefore, to gain resolution on the genuine function of histone marks, approaches and tools for their direct edition have been developed^11^. Manipulation of histone residues allowed revealing the key role of H3 methylations in animal and plant cell differentiation and specific developmental programs^12–15^. In a prior study involving the editing of the H3K27 residue in *Arabidopsis thaliana*, we not only confirmed expected functions for the H3K27me3 mark but also discovered novel roles in cell fates, critical for tissue regeneration and plant architecture through stem tissue differentiation^16^. While such global approaches have provided valuable insights, they affect the entire epigenome simultaneously, making it challenging to pinpoint the direct effect of a specific mark on a target gene^11^. Hence, novel CRISPR-Cas derived tools have been developed for various model organisms, serving as a platform to tether an effector capable of modifying the expression or epigenetic marks at a precise genomic locus^11,17–19^. These tools harbour a catalytically inactive (referred to as “dead”) form of Cas9 (dCas9), lacking endonuclease activity but retaining the ability to bind a single guide RNA (sgRNA)^20^. Thus far, dCas9 epigenetic editing tools have been more extensively assessed in animal cell cultures, in the aim to deposit or remove DNA methylation, histone acetylation or methylation, albeit with mitigated degrees of success^21–25^. In plants, only a limited number of studies have implemented CRISPR dCas9-based tools to manipulate epigenetic marks. These studies focused on editing DNA methylation^26,27,28^, acetylation at H3K27^2930^, and methylation at H3K4^30^ and H3K9^30,31^, primarily analysing molecular effects on the epigenetic mark and gene expression, without delving into the developmental consequences.

Here we present a novel approach utilising the CRISPR dCas9 SunTag system, to manipulate for the first time the repressive H3K27me3 mark at the organ boundary *CUP-SHAPED COTYLEDON 3 (CUC3)* gene. The rationale behind selecting this specific mark-gene pair is that H3K27me3 was reported to be a major determinant of tissue-specific expression patterns at the plant shoot apex^32^, where the *CUC3* gene is differentially expressed, delimitating the boundary between the shoot apical meristem and the organ primordia^33^ (Figure 1A). For this purpose, the Jumonji C-domain (JMJC) domain of the Arabidopsis JMJ13 demethylase^34^ (Figure 1B) was integrated into the dCas9 SunTag system^26^, allowing the recruitment of several effectors per locus, facilitated by an epitope-antibody amplification mechanism (Figure 1A; Figure S1).

**Figure 1.**
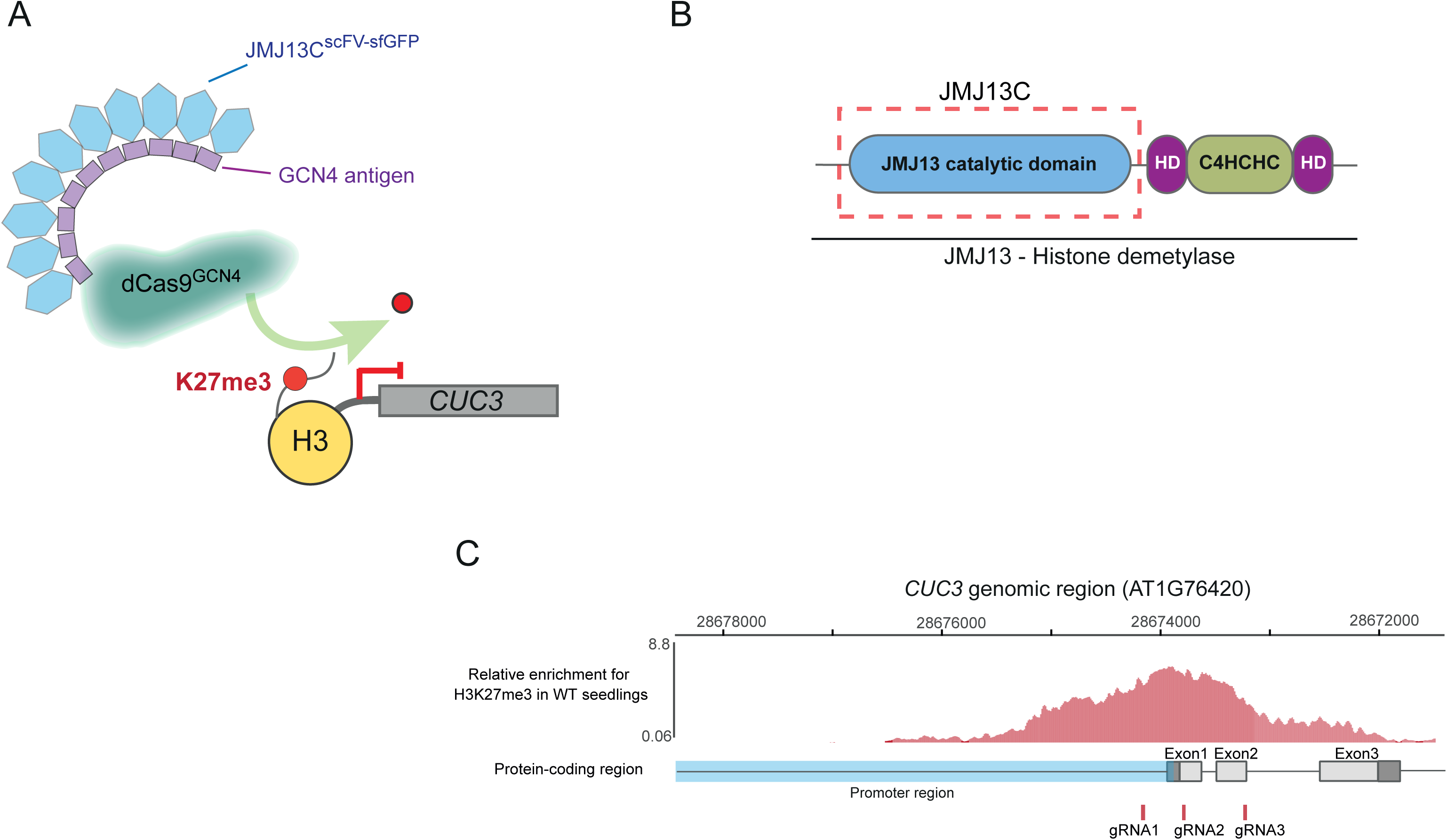
The dCas9-JMJ13^CUC3^ histone modification editor, a new tool designed to specifically remove H3K27me3 at *CUC3*. **(A)** Schematic representation of the chromatin editing approach for targeted removal (green arrow) of the repressive histone modification H3K27me3 (depicted as a red dot on the H3 histone tail) from the 5’ part of the *CUC3* gene region using the dCas9-based tool with the Sun-Tag amplification system. The dCas9^GCN4^ construct can recruit up to ten copies of the chromatin modifying module JMJ13C^scFV-sfGFP^ to target the *CUC3* regions via specific gRNAs. Violet bars: GCN4 antigen, present in 10 repeats; blue hexagons: JMJ13 C-terminal domain fused to the anti-GCN4 scFV (single-chain variable fragment) and sfGFP (Superfolder GFP). **(B)** Schematic representation of the Arabidopsis JMJ13 protein structure, containing the catalytic domain JMJC and the C4HCHC-type zinc finger domain^40^. The red dash-lined box outlines the protein region selected for use in this study. **(C)** Representation of the *CUC3* genomic region (AT1G76420), with the blue outline marking the promoter and the grey rectangles indicating the exons (dark grey delineates the 5’UTR and 3’UTR). The enrichment in H3K27me3 at this locus is illustrated by the red highlighted area^55^. Red lines below the *CUC3* genomic region indicate positions of guide RNAs designed in this study.

### Design and production of the dCas9-JMJ13^CUC^^3^ tool to manipulate the H3K27me3 mark at the CUC3 developmental gene

Several JMJ domain proteins in plants are known to act as histone demethylases^35–38^, with three of them specifically targeting H3K27^34,39–41^. Arabidopsis JMJ13, in particular, has been reported to contribute to photoperiod-dependent flowering regulation and self-fertility through the removal of histone methylation with high specificity towards repressive H3K27me3^38,40,41^. Based on the reported structure of JMJ13, we selected and cloned the JMJC catalytic domain of Arabidopsis JMJ13 to be incorporated into the CRISPR-dCas9 system, with the aim of precisely removing H3K27me3 at the selected region. The dCas9 SunTag amplification system was chosen based on its successful application in previous reports for DNA methylation editing in plants^26–28^.

*CUP SHAPED COTYLEDON3 (CUC3)* gene encodes a NAC domain family transcription factor that (along with *CUC1* and *CUC2*) plays a pivotal role in shoot meristem initiation and maintenance, organ initiation and separation, leaf shape, and positioning of the carpel margin meristems^33,42–45,46,47,48^. The expression of *CUC* genes is regulated through multiple pathways, including transcriptional control and post-transcriptional regulation by miRNAs of the miR164 family for *CUC1* and *CUC2* ^44,49–53^. The expression of *CUC3*, that lacks the miRNA target site, is positively regulated by *CUC2*^44,49,54^ . In addition, the *CUC3* gene region exhibits an enrichment in the repressive epigenetic mark H3K27me3 in leaf tissues as compared to shoot meristems^32^, indicating the contribution of epigenetic mechanisms to the regulation of its expression (Figure 1C). We thus hypothesised that the targeted removal of the repressive H3K27me3 at the *CUC3* region may help to better understand the contribution of this epigenetic modification to gene expression regulation and serve as proof of concept for the editing of this chromatin mark.

We designed three sgRNAs, based on available data for H3K27me3 enrichment in Arabidopsis seedlings^32,55^, to bring the dCas9-JMJ13 activity to the *CUC3* genomic region (Figure 1C). These sgRNAs were designed to target the promoter and proximal parts of the gene. Specifically, gRNA1 is positioned within the promoter region, gRNA2 near the transcription start site (TSS), and gRNA3 within the first exon of *CUC3*.

We hereinafter refer to the epigenetic editing tool developed in this study as the dCas9-JMJ13^CUC3^ tool. To assess its impact, the reporter line *pCUC3::CFP*^44^ was selected as the recipient for the dCas9-JMJ13^CUC3^ editing tool. This choice facilitates the monitoring of transcription from the *CUC3* promoter as well as expression from the *CUC3* endogenous locus.

Several independent transgenic lines were produced, carrying constructs with or without the JMJ13 catalytic domain, thereafter referred to as SunTagJMJ13gCUC3 and SunTag_gCUC3, respectively (SunTagJMJ13gCUC3: 41 primary transformants, 10 analysed lines at the T2 generation, among which 4 were included in the study for analyses on the T3 and T4 generations; SunTag_gCUC3: 90 primary transformants, 10 analysed lines at the T2 generation, among which 2 were included in the study for analyses on the T3 and T4 generations). The effects of the dCas9-JMJ13 tool on developmental features and target gene expression were deduced from analyses on the SunTagJMJ13gCUC3 plants in comparison to SunTag_gCUC3 and untransformed *pCUC3::CFP* (thereafter referred to as WT) plants.

### The dCas9-JMJ13^CUC^^3^ tool induces developmental phenotypes characteristic of *CUC3* ectopic expression

Under long-day conditions, the plants of SunTagJMJ13gCUC3 lines displayed lower growth rates as compared to WT and SunTag_gCUC3 plants (Figure 2A). Specifically, for the four analysed SunTagJMJ13gCUC3 independent lines, the areas of the rosette leaves are significantly smaller than those of the two independent SunTag_gCUC3 control lines (Figure 2B, Figure S2). Additionally, the rosette leaves of dCas9-JMJ13 plants have an overall lower length-to-width aspect ratio than control plants (Figure 2C). These smaller rosette and rounder leaf phenotypes are similar to those, earlier reported, of plants conditionally over-expressing *CUC3* (*p35S::CUC3-GR* transgenic lines), and correspond well to the known functions of the CUC3 transcription factor as a growth repressor^42,56^.

**Figure 2.**
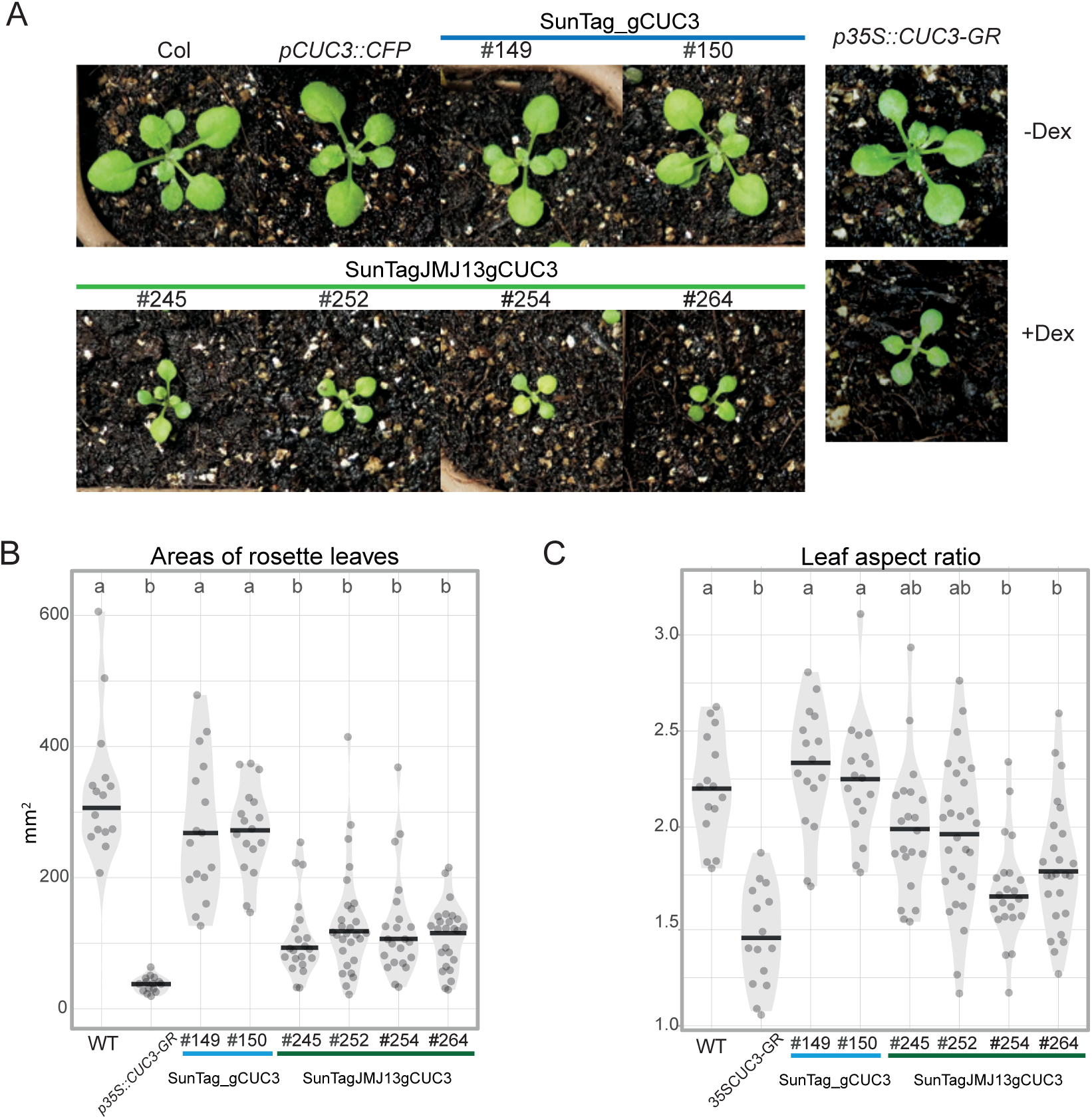
The dCas9-JMJ13^CUC3^ tool induces rosette phenotypes associated with *CUC3* ectopic expression. **(A)** Representative images of 16-day-old plantlets grown at 21°C under long-day conditions. The upper panel features (from left to right) plants from the wild-type Col ecotype, the *pCUC3::CFP* (WT) line, and two independent transgenic lines containing the dCas9 construct without the JMJ13 catalytic domain (SunTag_gCUC3). The lower panel features plants from four independent transgenic lines harbouring the dCas9 construct with JMJ13 catalytic domain (SunTagJMJ13gCUC3). The right panel displays images of plants from the inducible *p35S::CUC3-GR* line, grown on soil, in absence (-Dex) or presence (+Dex) of dexamethasone. Diagrams showing **(B)** the average surface (mm^2^) and **(C)** aspect ratio of leaves (length:width) for each genotype mentioned in (A). Sample size: *n* = 16, 15, 16, 17, 21, 28, 22 and 25 for WT, p35S::CUC3-GR, #149 and #150 (SunTag_gCUC3), #245, #252, #254 and #264 (SunTagJMJ13gCUC3), respectively. Black lines represent medians and dots values of individual samples. Letters indicate significant differences (Tukey pairwise comparison test, P<0.05).

SunTagJMJ13gCUC3 adult plants also display noticeable developmental phenotypes. Notably, we detected the splits of shoot apical meristems in all four SunTagJMJ13gCUC3 lines, occurring with various frequencies (between 28% for line #245 and 43% for line #252) (Figure S3 A, B). After final elongation, the SunTagJMJ13gCUC3 plants, on average, initiated a higher number of stems from rosette and displayed a trend toward shorter overall inflorescence stem length (Figure S3 C, D). While these traits presented some variability within plants of the same line and between independent lines, they consistently displayed a trend significantly different from the control lines (WT and SunTag_gCUC3). As a matter of fact, the ectopic expression of *CUC* genes has also been associated with an increase in branching^52^.

Together, these observations provide good indication that the dCas9-JMJ13^CUC3^ tool leads to ectopic de-repression of *CUC3*, likely as a result of the intended decrease in the repressive H3K27me3 mark. To verify this, we conducted further experiments on three of the SunTagJMJ13gCUC3 lines in comparison to SunTag_gCUC3 lines and a WT control, all in the *pCUC3::CFP* background.

### dCas9-JMJ13^CUC^^3^ leads to activation of *CUC3* transcription, within its expression territory and ectopically

We further analysed the effects of *dCas9-JMJ13^CUC^*^3^ on its target transcription and expression in seedlings, using two distinct approaches. Firstly, CFP fluorescent signal produced from the *pCUC3::CFP* construct was used for analysis of transcription from the *pCUC3* promoter. CFP signals were visualised by epifluorescence microscopy on 10-day-old seedlings from all test and control lines, and quantified from pictures taken on individual samples (Figure 3A, B, Figure S4). While heterogeneity in signal intensity was present among the seedlings within each line, quantification of an overall area covered by fluorescent signal showed that it was significantly more intense, as well as larger in seedlings of the SunTagJMJ13gCUC3 lines compared to the SunTag_gCUC3 and WT lines. This indicates both a stronger transcriptional activity from the *pCUC3* promoter, but also a broader domain of expression within the seedling tissue. Secondly, to assess *CUC3* expression from the endogenous locus, we employed RT-qPCR, comparing rosettes from the SunTagJMJ13gCUC3 and SunTag_gCUC3 lines. The level of *CUC3* mRNA was increased from 2 to 7-fold depending on the plant and line. While the *dCas9-JMJ13^CUC^*^3^ construction has a significant overall effect on *CUC3* expression, heterogeneity in response between cells within a same tissue may account for the differences observed between plants of a same line, as indicated by the *in situ* CFP fluorescence imaging (Figure S4). Yet, together, relative expression trends observed by RT-qPCR among lines were in agreement with results of the *pCUC3::CFP* fluorescence analyses (Figure 3, Figure S4), and indicate a dCas9-JMJ13-induced de-repression of transcription at the *pCUC3* promoter and at the *CUC3* locus.

**Figure 3.**
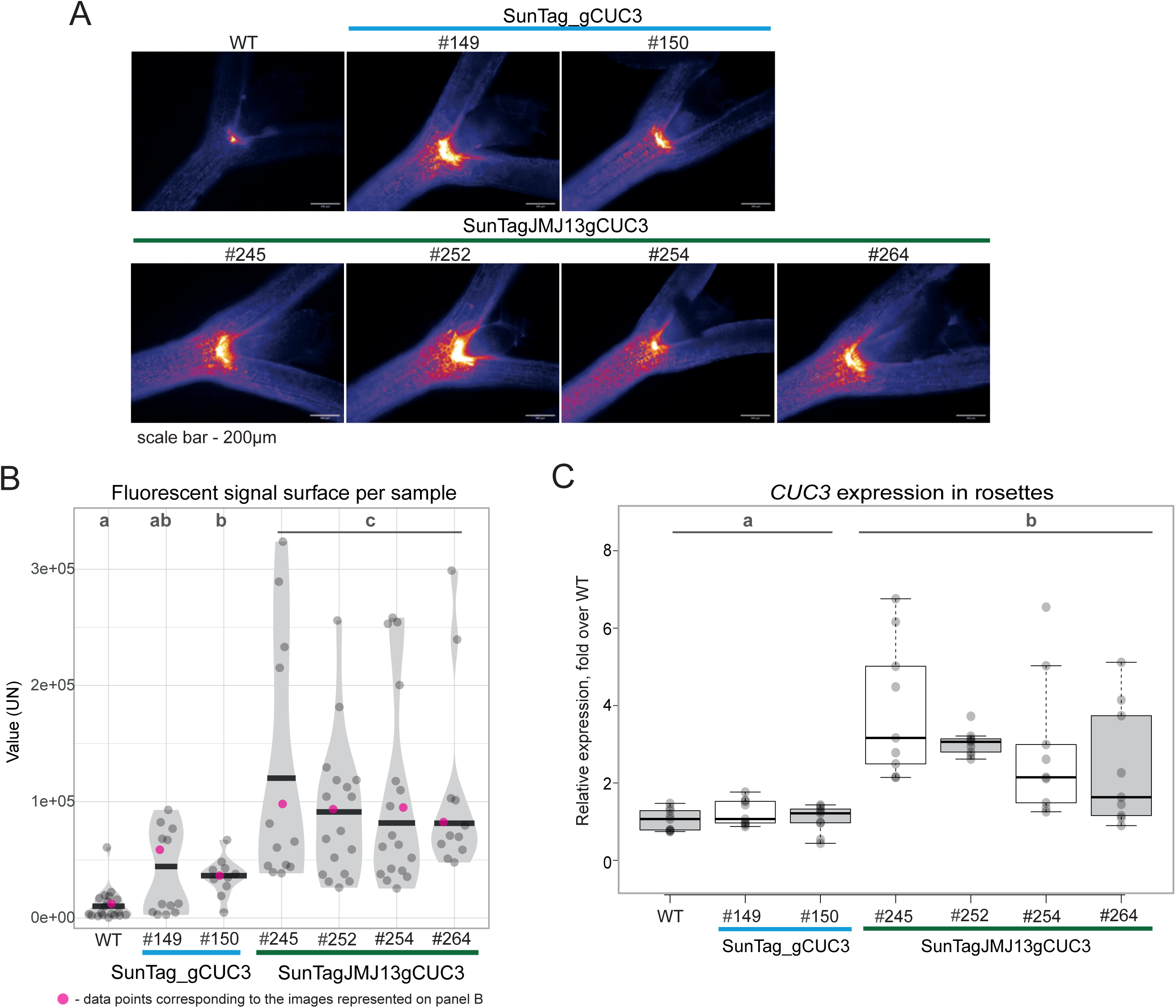
Transcription from the *pCUC3* promoter and expression of *CUC3* are induced in SunTagJMJ13gCUC3 lines. **(A)** Representative fluorescence microscopy images of the 10-day-old seedlings visualising the CFP reporter expressed from the *CUC3* promoter (*pCUC3::CFP*). The upper panel displays the plants of wild type and two independent transgenic lines that contain the dCas9 construct without the JMJ13 catalytic domain (SunTag_gCUC3). The lower panel displays the representative plant images of four independent transgenic lines that contain the dCas9 construct with JMJ13 catalytic domain (SunTagJMJ13gCUC3). Scale bars: 200µm. **(B)** Violin plots illustrating the quantification of the fluorescent signal surfaces on individual microscopy samples (seedlings). Sample size: *n* = 19, 13, 11, 13, 18, 18 and 12 for WT, #149 and #150 (SunTag_gCUC3), #245, #252, #254 and #264 (SunTagJMJ13gCUC3), respectively. Black lines represent the median and the dots represent the values of individual samples; the samples are assembled in statistical groups by the Tukey pairwise comparison test. **(C)** Boxplots representing the relative expression of *CUC3* in the seedlings of the control and test lines. *TUBULIN* was used as a reference gene for normalisation. Black lines represent the median and the dots represent values scored for individual seedlings. Letters indicate significant differences (Tukey pairwise comparison test, P<0.05).

### Decreased level of H3K27me3 at CUC3 correlates with its transcriptional reactivation

Finally, to assess if the dCas9-JMJ13^CUC3^-induced changes in *CUC3* expression were due to an expected, significant decrease in the H3K27me3 mark, we analysed its abundance at the *CUC3* locus in seedlings, for all SunTagJMJ13gCUC3 transgenic lines that displayed robust phenotypes and effects on target gene expression. Using ChIP-qPCR, we detected that the amount of H3K27me3, reported to the amount of H3, was indeed lower in the SunTagJMJ13gCUC3 lines as compared to the control lines (WT and SunTag_gCUC3). This effect was the strongest within the first exon of *CUC3*, with a 5 to 10-fold decrease in H3K27me3 abundance, while the mark amount was reduced of 3 to 5 folds in the second exon (Figure 4, Figure S5). Interestingly, according to ChIP-seq data, the first exon is the region of *CUC3* locus where H3K27me3 is most abundant (Figure 1C). Importantly, no significant decrease in H3K27me3 was detected in the SunTag_gCUC3 control, supporting the functionality (H3K27me3 demethylase effect) of the chosen JMJ13 catalytic domain when fused to the dCas9 SunTag system.

**Figure 4.**
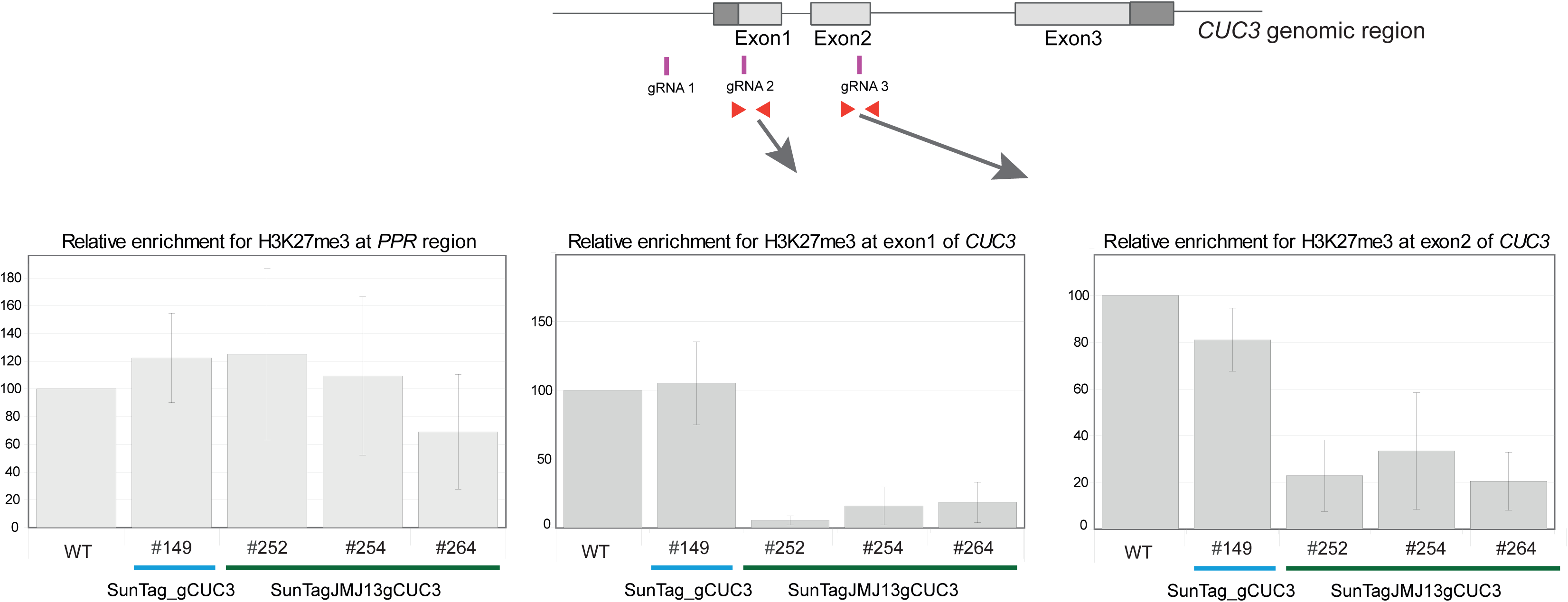
The dCas9-JMJ13^CUC3^ tool induces reduction in the H3K27me3 mark abundance at the *CUC3* gene region, in SunTagJMJ13gCUC3 lines. Histograms illustrating the relative enrichment for H3K27me3 at two regions of *CUC3*, depicted by the schematic drawing on the top, as detected by ChIP-qPCR. The *PPR* (AT5G55840) gene region was used as a negative control. The relative H3K27me3 enrichment was calculated as a fold change between the percentage of input enrichment obtained after immunoprecipitation with the anti-H3K27me3 antibody, over that obtained with the anti-H3 antibody for the corresponding samples, and is represented relative to WT (set to 100). Each histogram bar corresponds to the mean value (the error bars indicates the standard deviation), calculated from of 3 biological repeats (for each repeat, the PCR quantification was performed with 3 technical replicates). The individual results of the 3 independent ChIP experiments can be visualised in Figure S5.

In conclusion, we have reported here the use of a CRISPR dCas9-based system employing the JmJ13 catalytic domain to selectively remove the repressive H3K27me3 mark and thereby manipulate transcription from the organ frontier gene *CUC3* in Arabidopsis.

Our results show that the inflicted decrease in the repressive epigenetic mark at targeted regions results in the de-repression of *CUC3* in plant tissues and is associated with developmental phenotypes. This comprehensive dataset provides a proof-of-concept, seamlessly bridging molecular insights to developmental evidence. It thus validates a valuable approach to resolve the roles of individual histone marks in the regulation of chromatin structure and transcription dynamics in plants, with an ultimate readout on cell fate.

With our characterisation of dCas9-JMJ13^CUC^^3^, precise chromatin edition tools proved instrumental in assessing if chromatin marks can be primary determinants of gene expression and cell differentiation. They likely could be pushed further toward inducible systems for more precise post-perturbation analyses, thereby allowing to explore changes in the nucleus and chromatin structure, cross-talks between epigenetic marks, and effect on transcription kinetics.

## Materials and Methods

### Cloning and generation of transgenic lines

sgRNA design was performed using the CHOPCHOP tool (http://chopchop.cbu.uib.no/, Repair profile prediction^57^ combined with Cas-Offinder (http://www.rgenome.net/cas-offinder/) and TAIR blast tools for verification of off-target effects. The qRNA cassette was custom-synthesised by GenScript (www.genscript.com) and inserted into into SunTag dCas9 plasmid (Addgene Plasmid #117168) using the KpnI and MauBI restriction enzymes (Thermo Scientific™, ER0522 and ER2081 respectfully). The JMJ13 catalytic domain was amplified with the primers listed in Table S1 and cloned into SunTag dCas9 plasmid using the BsiWI restriction enzyme (Thermo Scientific™, ER0851). The final construct allows to produce (i) a dCas9 fusion to 10 copies of the short epitope GCN4, (ii) a superfolderGFP-JMJ13 effector domain combination fused to a single-chain variable fragment - scFV-antibody directed against GCN4, and (iii) three sgRNA complementary to *CUC3* genomic sequence (Figure S1).

### Plant culture and phenotyping

All plants were cultured in growth chambers, in long-day conditions, 16 h/8 h light/dark period, at 21°C. For the selection of transgenic lines, the seeds of transformed plants were germinated and grown for 10 days on Murashige-Skoog (MS) plates containing Hygromycine B (Merck H3274). Resistant plants were transferred to soil and genotyped with the primers listed in Table S1. Lines with a single insertion locus were brought to the T3 generation for further characterization. The procedures for quantitative phenotype characterisation were performed on plants of T3 and T4 generations grown.

Detection of the CFP expression in the tissues was performed on 10-day-old MS plate grown seedlings. Images were acquired using the Zeiss Imager.M2 microscope (20× and 40× objective) with the Axiocam 503.

The size of rosettes was assessed from images of 15-day-old plants using the FIJI software ^58^, by drawing circles that touched the extremities of 3 rosette leaves on each plant. The areas and aspect ratio of rosette leaves were measured by outlining the contour of the third true leaf on individual plants within the population.

The inflorescence stem length and quantity of side branches were quantified on plants with fully elongated main stems after all flowers were opened.

### Plot preparation and statistical analysis

Plots of all presented data sets were prepared using the Rstudio software (*RStudio Team (2020),* http://www.rstudio.com/). The Tukey’s range test was used to make the pairwise comparisons of means from independent samples.

### Gene expression analyses

Expression of the transgene and *CUC3* in the generated lines were verified by RT-qPCR. RNA was extracted from rosette leaves of 15-day-old plants and purified using the Qiagen RNeasy Plant Mini Kit (Cat. No. / ID: 74904). After DNase treatment (ezDNase SuperScript IV VILO, ThermoFisher, Cat. No. 11756050), first strand cDNA synthesis was performed from 2µg of total RNA using SuperScript IV VILO (ThermoFisher, Cat. No. 11756050). Relative transcript abundance was measured using the SYBR Green Master Mix (POWER SYBR GREEN PCR, Thermo Fisher Scientific, 10658255) on a CFX Connect BioRad Real-Time PCR System. Gene-specific primers used for amplification are listed in Table S1.

### Chromatin Immunoprecipitation

Chromatin fraction was isolated from 10-day-old seedlings following the procedure described in^55^. The antibodies used were anti-trimethyl-H3K27 (07-449 Millipore) and anti-H3 (AS10710 Agrisera). Reverse-cross-linked samples were purified using the Qiagen Reaction Minelute Kit (#T1030L) with an elution volume of 20μl. The procedure was carried out on samples collected and prepared from 3 independently grown plant populations. Immuno-precipitation was performed on chromatin extracts, using either the anti-H3K27me3 antibody or the anti-H3 antibody. The ChIP-qPCR for selected target regions was performed as described above for the RT-qPCR, with 3 technical replicates, using the primers listed in Table S1. The H3K27me3 enrichment was calculated relatively to that obtained after immunoprecipitation with the anti-H3 antibody for each corresponding sample.

## Acknowledgments

We thank Anne-Marie Boisson, Dila Cetin, Adrien Galeone, Emilien Krempf, Alizée Musso, and Mirko de Vivo for help with plant culture, selection and characterisation of transgenic lines.

## Author contributions

Conceptualization, C.C.C. and K.F.; methodology, C.C.C, K.F. and A.B.; formal analysis, C.C.C. and K.F.; investigation, K.F., M.L.M. and C.C.C.; writing – original draft, K.F. and C.C.C.; writing – review & editing, C.C.C. with help of A.B. and K.F.; funding acquisition, C.C.C. and A.B.

## Funding

This work was supported by the Agence Nationale de la Recherche (ANR-18-CE20-0011-01, PRC project REWIRE to C.C.C. and A.B.) and the Grenoble Alliance for Cell and Structural Biology (ANR-10-LABX-49-01).

## Supplementary information

**Supplementary Figure S1.**
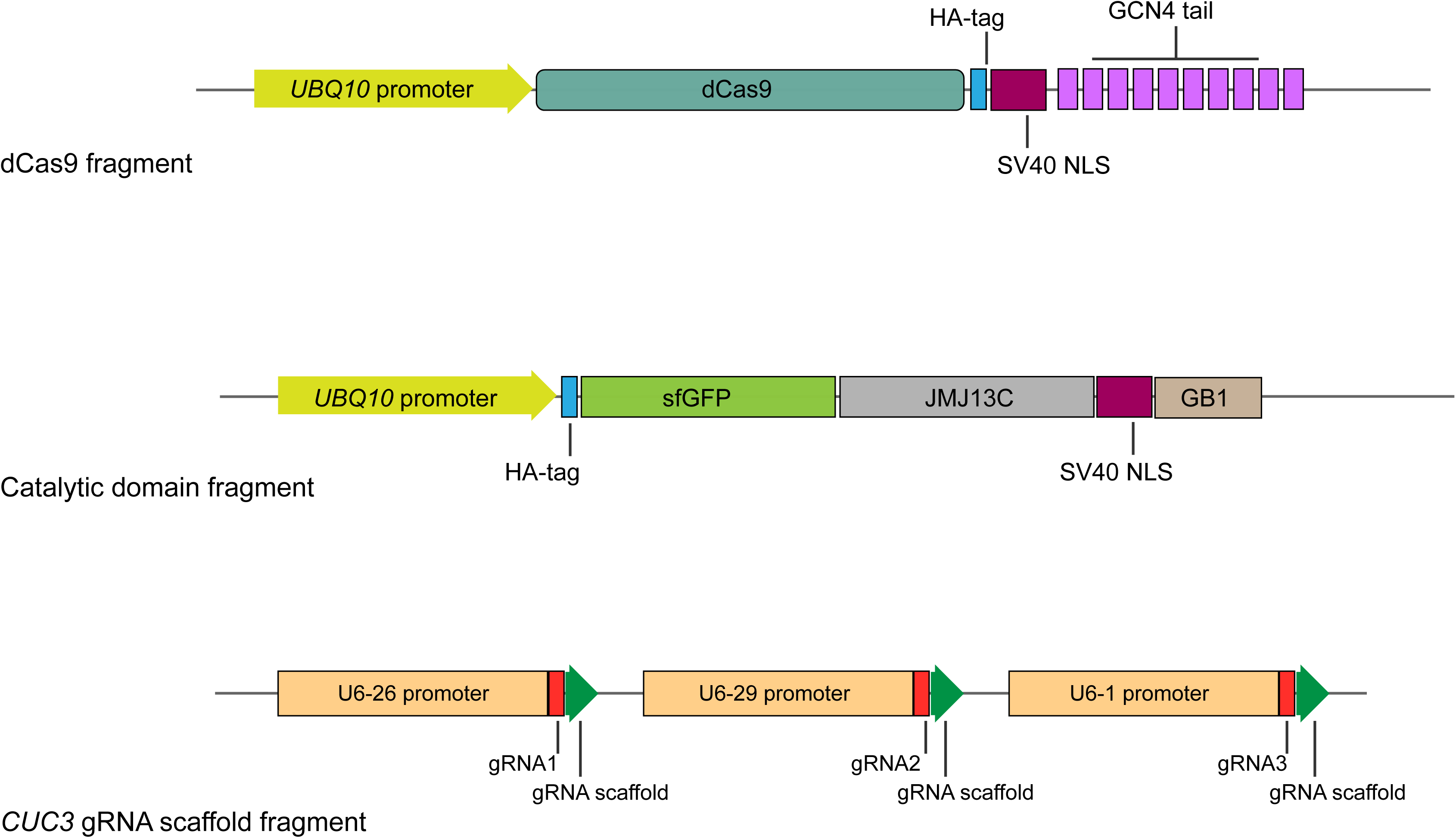
Schematic illustrating the three modules of the SunTag construct. The dCas9^GCN4^ module consists of dCas9 fused to a tail made of 10 copies of the GCN4 epitope and a triple SV40 NLS, whose expression is controlled by the UBQ10 promoter. The JMJ13C^scFv-sfGFP^ module consists in the Catalytic domain of JMJ13 fused to scFv-sfGFP and a GB1-REX NLS (NLS sequences present in the SunTag construct reported in Papikian *et al*., 2019), whose expression is also controlled by the UBQ10 promoter. The gRNA module consists of three sequential expression cassettes with gRNAs whose expression is controlled by independent U6 promoters (U6-26, U6-29 and U6-1).

**Supplementary Figure S2.**
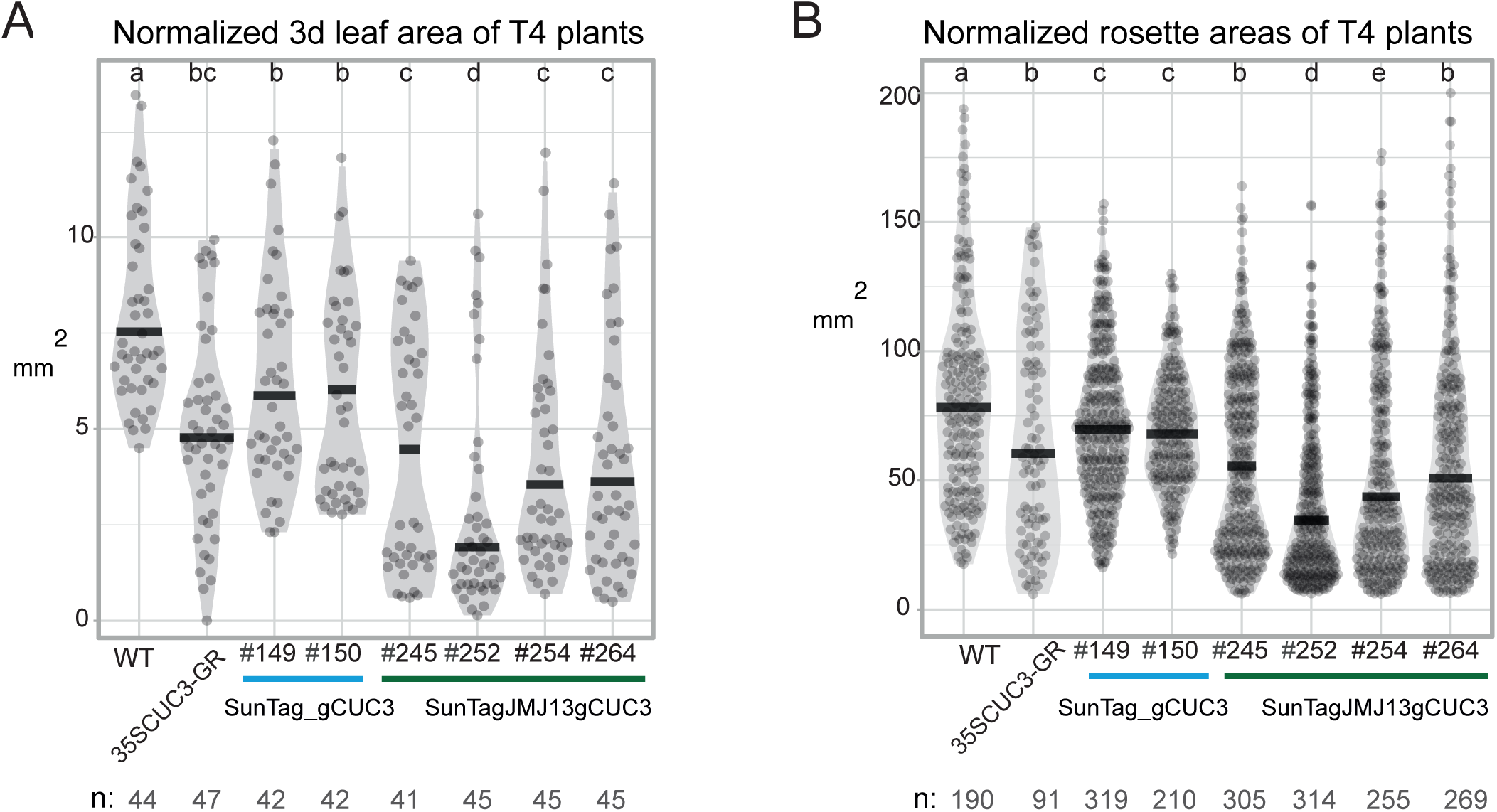
The dCas9-JMJ13CUC3 tool induces rosette phenotypes associated with *CUC3* ectopic expression. Diagrams showing the average leaf (A) and rosette (B) surface areas (mm^2^) for the plants of WT, *p35S::CUC3-GR,* SunTag_gCUC3 and SunTagJMJ13gCUC3 genotypes. The surface areas of the third leaf were measured plants from two independent T4 populations with the total sample size: *n* = 44, 47, 42, 42, 41, 45, 45 and 45 for WT, p35S::CUC3-GR, #149 and #150 (SunTag_gCUC3), #245, #252, #254 and #264 (SunTagJMJ13gCUC3), respectively. The rosette area measurements were acquired on plants from two independent T4 populations with the total sample size: *n* = 190, 91, 319, 210, 305, 314, 255 and 269 for WT, p35S::CUC3-GR, #149 and #150 (SunTag_gCUC3), #245, #252, #254 and #264 (SunTagJMJ13gCUC3), respectively. Black lines represent medians and dots values of individual samples. Letters indicate significant differences (Tukey pairwise comparison test, P<0.05).

**Supplementary Figure S3.**
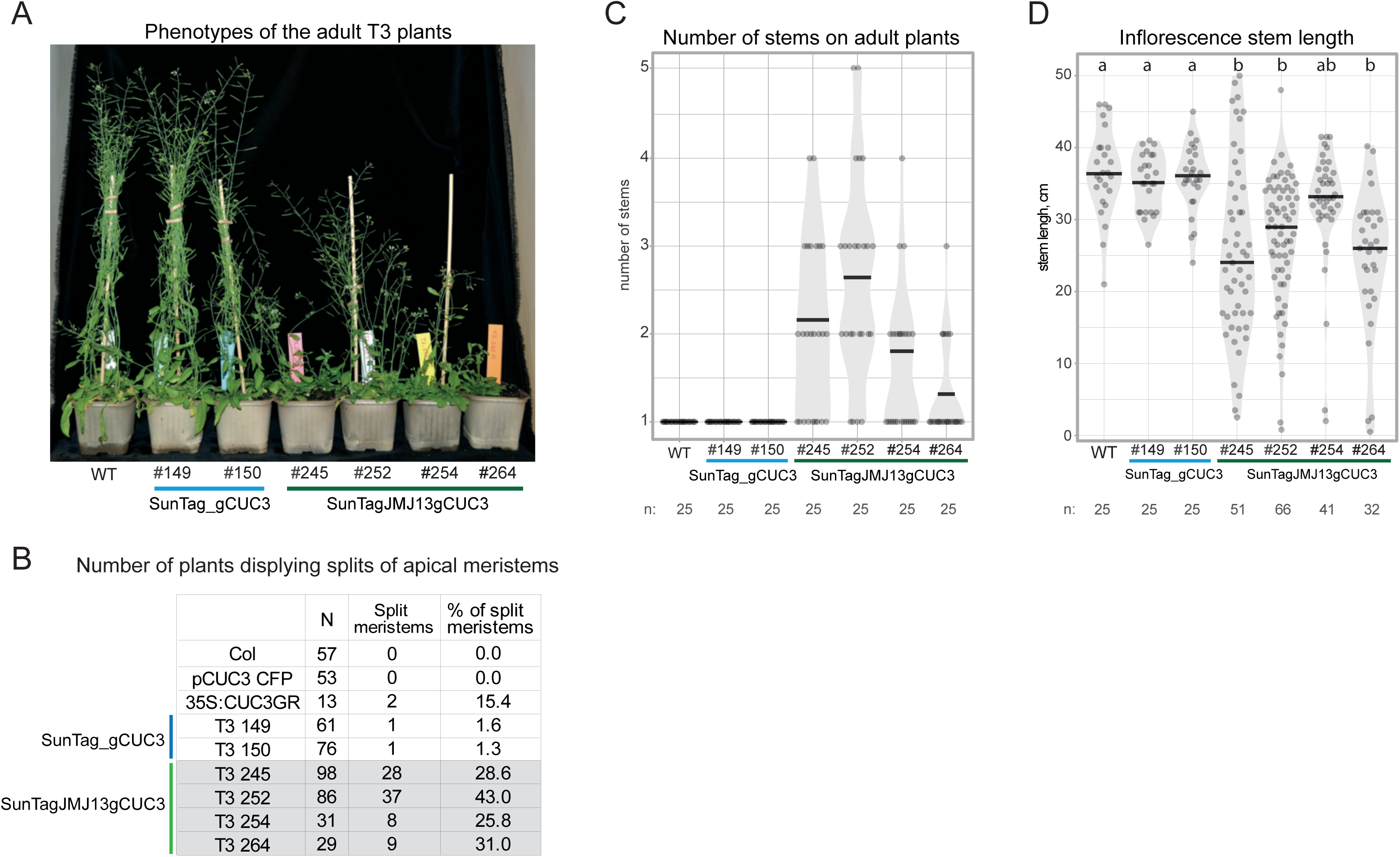
Adult plant phenotypes, associated with the dCas9-JMJ13CUC3 tool and *CUC3* ectopic expression. (A) Representative image of the adult plants of (from left to right) WT line, two independent transgenic lines containing the dCas9 construct without the JMJ13 catalytic domain (SunTag_gCUC3) and four independent transgenic lines caring the dCas9 construct with JMJ13 catalytic domain (SunTagJMJ13gCUC3). All pictured plants belong to the simultaneously sawn populations, grown at 21°C under long-day conditions. (B) Table, illustrating the average number of plants displaying splits of apical meristems within three independently grown populations. (C) Diagram displaying the average number of inflorescence stems on the plants from each of the genotypes, mentioned in (A) with the total sample size of *n* = 25 for all the genotypes. (D) Diagram illustrating the average maximal inflorescence stem length for the plants from each of the genotypes, mentioned in (A) with the sample size of *n* = 25, 25, 25, 51, 66, 41, and 32 for WT, #149 and #150 (SunTag_gCUC3), #245, #252, #254 and #264 (SunTagJMJ13gCUC3), respectively. All phenotype quantification measurements for C and D were acquired on plants from two independent T4 populations. Black lines represent medians and dots values of individual samples. Letters indicate significant differences (Tukey pairwise comparison test, P<0.05).

**Supplementary Figure S4.**
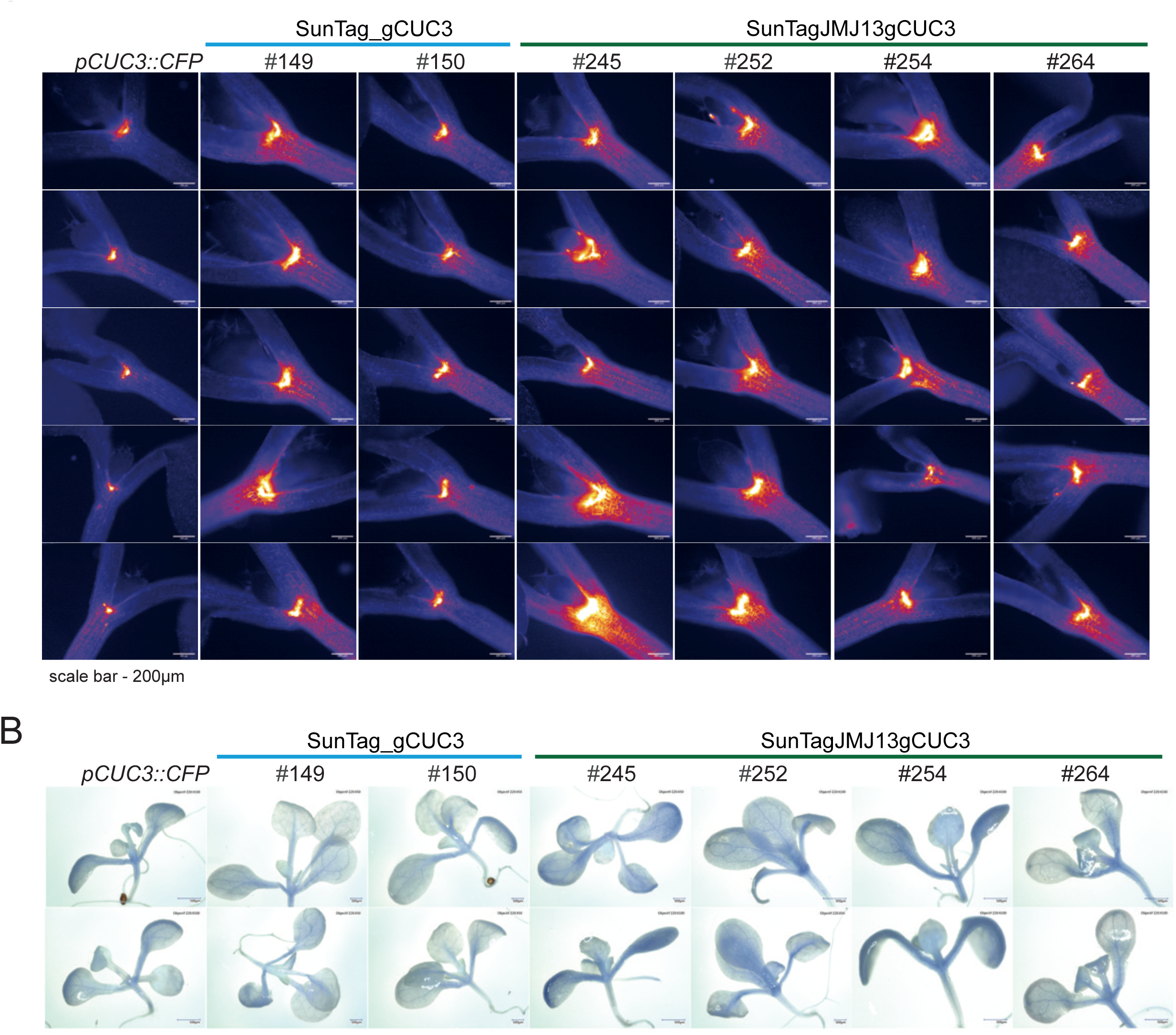
Representative fluorescence microscopy images of the 10-day-old seedlings, visualizing the CFP reporter expressed from the *CUC3* promoter (*pCUC3::CFP*). The columns from left to right display the plants of wild type and two independent transgenic lines that contain the dCas9 construct without the JMJ13 catalytic domain (SunTag_gCUC3) followed by four independent transgenic lines that contain the dCas9 construct with JMJ13 catalytic domain (SunTagJMJ13gCUC3). Scale bars: 200µm.

**Supplementary Figure S5.**
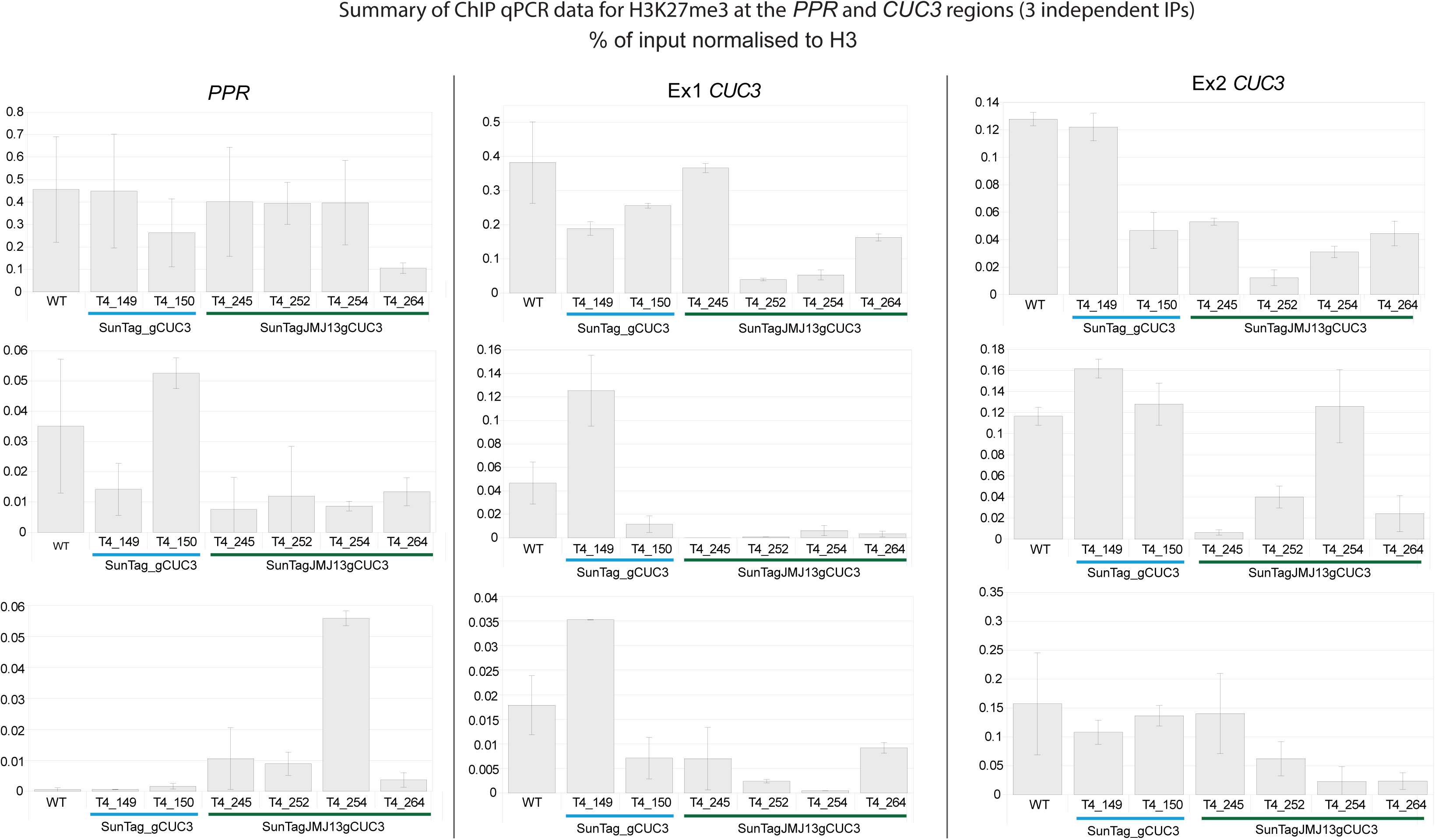
The dCas9-JMJ13CUC3 tool induces reduction in the H3K27me3 mark abundance at the *CUC3* gene region, in SunTagJMJ13gCUC3 lines. Histograms illustrating the relative enrichment for H3K27me3 at two regions of *CUC3*, depicted by the schematic drawing on Figure 4, as detected by ChIP-qPCR. The *PPR* (AT5G55840) gene region was used as a negative control. The relative H3K27me3 enrichment was calculated as a fold change between the percentage of input enrichment obtained after immunoprecipitation with the anti-H3K27me3 antibody, over that obtained with the anti-H3 antibody for the corresponding samples, and is represented relative to WT (set to 100). The individual results of the 3 independent ChIP experiments are presented in 3 independent graphs organised in column, with the mean values (and standard deviation) for each histogram calculated from 3 technical replicates. Only one line (#245) out of the four tested did not display consistent changes between replicates.

**Supplementary Table S1.** Information on primers used in this study.

## Notes

### Competing Interest Statement

The authors have declared no competing interest.

